# Genetic dissection of MAPK-mediated complex traits across *S. cerevisiae*

**DOI:** 10.1101/004895

**Authors:** Sebastian Treusch, Frank W. Albert, Joshua S. Bloom, Iulia E. Kotenko, Leonid Kruglyak

## Abstract

Signaling pathways enable cells to sense and respond to their environment. Many cellular signaling strategies are conserved from fungi to humans, yet their activity and phenotypic consequences can vary extensively among individuals within a species. A systematic assessment of the impact of naturally occurring genetic variation on signaling pathways remains to be conducted. In *S. cerevisiae*, both response and resistance to stressors that activate signaling pathways differ between diverse isolates. Here, we present a quantitative trait locus (QTL) mapping approach that enables us to identify genetic variants underlying such phenotypic differences across the genetic and phenotypic diversity of *S. cerevisiae*. Using a Round-robin cross between twelve diverse strains, we determined the genetic architectures of phenotypes critically dependent on MAPK signaling cascades. Genetic variants identified fell within MAPK signaling networks themselves as well as other interconnected signaling pathways, illustrating how genetic variation can shape the phenotypic output of highly conserved signaling cascades.

## Highlights

- Quantitative trait mapping using large mapping populations from natural isolates
- QTL mapping for MAPK-mediated traits
- Identification of genetic variants that influence cellular signaling pathways

## Introduction

Cellular survival is dependent on the ability to sense and respond to changing environmental conditions. Mitogen-activated protein kinase (MAPK) signaling cascades are ubiquitous in eukaryotic organisms and enable them to react to extracellular stimuli [1]. MAPK cascades are composed of three sequentially acting kinases [2], which upon sensing an extracellular stimulus trigger a cellular response by activating transcription factors and other regulatory proteins [2,3]. The yeast *Saccharomyces cerevisiae* has long been used as a eukaryotic model system to elucidate the principles of these signaling pathways, as many of the core signaling proteins are conserved from yeast to human [3].

In yeast, MAPK pathways facilitate response to environmental cues such as the presence of mating pheromones and stresses such as cell wall damage and high osmolarity [4]. Adaptation to high osmolarity is conducted by the HOG (high osmolarity glycerol) MAPK pathway. The human homolog of the MAPK Hog1, p38α, not only mediates the response to hyperosmolarity as well [4], but also plays key roles in inflammation and cancer [5,6]. Insults to the cell wall are sensed by the cell-wall integrity (CWI) pathway, which is anchored by the MAPK Slt2, homolog of the mammalian MAPK7 [7]. Response and resistance to MAPK-activating stress conditions are highly variable among different yeast isolates [8,9,10]. While genes of the core MAPK cascades are highly conserved across species, upstream regulatory components, such as stress sensors, and downstream targets, such as transcription factors, exhibit high levels of divergence [11,12]. Yet, it is unknown how genetic differences in such elements of the MAPK pathways contribute to the phenotypic differences between isolates of the same species. Hence, it remains poorly understood how sequence variation among yeast isolates results in quantitative differences in MAPK-dependent phenotypes.

Yeast is an ideal model system for the study of quantitative traits. Quantitative trait locus (QTL) mapping studies in *S. cerevisiae* have characterized the genetic architecture of complex traits, including global gene expression and resistance to small molecules [13,14,15,16], as well as shed light on the sources of “missing heritability” [17]. We have previously developed a bulk-segregant-analysis approach (BSA), named X-QTL, which utilizes extremely large mapping populations for increased mapping power [18]. Such mapping populations are subjected to selection to enrich for alleles that contribute to a trait of interest. We have recently used this method to determine the genetic underpinnings of protein abundance variation [19]. The majority of mapping studies to date have interrogated a pair of strains at a time, with only some recent studies expanding to crosses among four parent strains [20,21]. Isolation of individual recombinant haploids, as used in linkage mapping studies, is very labor intensive, and current X-QTL protocols require extensive strain engineering. Consequently, mapping studies have queried a fraction of the *S. cerevisiae* species-wide genetic and phenotypic variation and have left much of the genetic architecture of quantitative traits across the species unexplored.

Here, we report an approach that enables implementation of X-QTL in crosses among diverse yeast isolates without extensive genetic modifications. This method takes advantage of plasmid-borne fluorescent markers to leverage the precision and high throughput of fluorescence-activated cell sorting (FACS) to rapidly generate large mapping populations of recombinant haploids. We dissected the genetic architectures of salt (sodium chloride) and caffeine tolerance in a set of strains capturing much of the species-wide genetic diversity. Salt tolerance is mediated by the HOG MAPK pathway, while exposure of yeast to caffeine results in the activation of the CWI MAPK pathway [7]. In addition to these two pathways, we examined the effect of the MAPK mating pathway on growth by comparing mapping populations of opposite mating type. We identified QTL that contain genes with various roles in MAPK signaling. Our results how genetic variation shapes MAPK-dependent traits. Finally, we illustrate how a round-robin cross design can be leveraged to identify causal genetic variants.

## Results

### Fluorescence-based isolation of mapping populations

Our original X-QTL approach relied on the expression of the auxotrophic marker *HIS3* under the control of the MATa-specific *STE2* promoter for the isolation of MATa recombinant haploids [22]. However, most isolates of *S. cerevisiae* are generally prototrophs, lacking the auxotrophic mutations required for the original X-QTL approach. In addition, auxotrophic mutations can have large phenotypic effects [14]. Furthermore, the use of the *STE2* promoter restricted X-QTL mapping populations to segregants of the MATa mating type. Here, we designed a novel approach that relies on mating-type specific expression of plasmid-borne fluorescent reporters, enabling the isolation of haploids of either mating type without the need to integrate the reporters into the genome (Figure 1A). We placed the fluorescent markers mCitrine and mCherry under the control of the *STE2* promoter and its MATα-specific equivalent, the *STE3* promoter, respectively [22]. We combined these two fluorescent markers in a single construct and created a set of plasmids with drug resistances that permit facile introduction into any strain background (see Methods for details).

**Figure 1:**
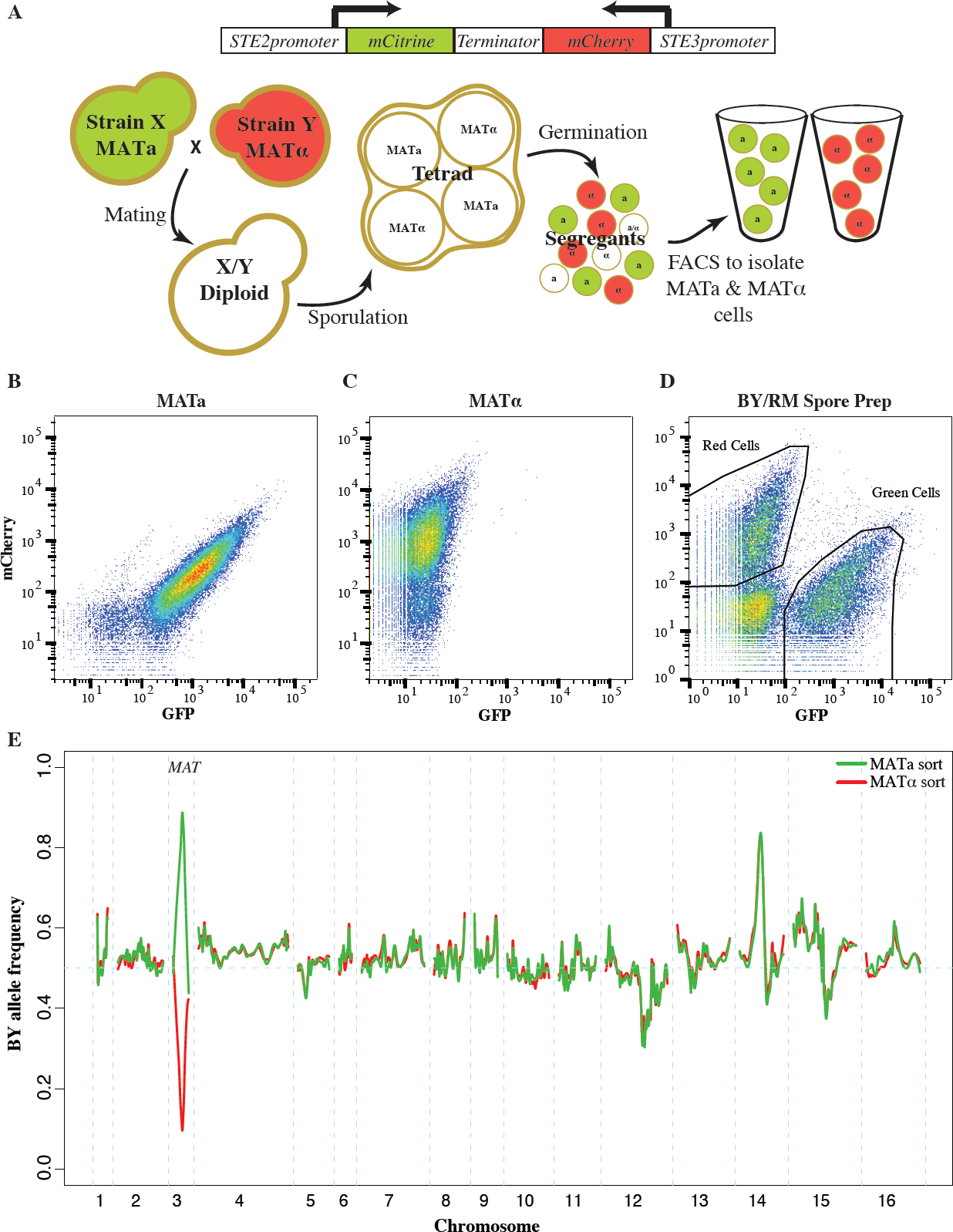
Fluorescent markers for the isolation of recombinant haploids. **(A)** Utilizing fluorescent mating type markers to generate large mapping populations. The fluorophores mCitrine and mCherry were placed under the control of the MATa-specific *STE2* and the MATα-specific *STE3* promoters, respectively, and combined into a single construct. Introduction of this construct into MATa cells results in green fluorescence, while MATα cells exhibit red fluorescence. Neither of the two reporters is active in diploid or tetrad cells, but after germination the markers are expressed and enable the isolation of MATa and MATα populations of recombinant haploids. **(B-E)** Validation of the fluoresent markers. Flow cytometry of **(B)** MATa cells and **(C)** MATα cells that carried a plasmid with the combined fluorescent mating type markers. MATa cells were green fluorescent, while MATα cells presented with red fluorescence. **(D)** A culture of BY/RM recombinant haploids with the mating type marker plasmid as assessed during FACS. Green MATa and red MATα cells were isolated based on their fluorescence. Gates shown represent approximate gates used during cell sorting. **(E)** The allele frequency spectrum across the genome within the isolated MATa and MATα cell populations was determined by sequencing. On chromosome three MATa populations were highly enriched for the MATa allele, while MATα populations were highly enriched for the MATα allele. Other large allele frequency skews correspond to previously identified BY/RM growth QTL.

As expected, the presence of this construct resulted in green fluorescence in MATa strains and red fluorescence in MATα strains (Figure 1B & C). Diploids carrying the marker construct were non-fluorescent, as neither promoter is active in diploid cells [22,23]. After sporulation and germination, haploid progeny express the fluorescent marker corresponding to their mating type. We tested this aspect of the reporter construct by introducing it into the diploid hybrid between the well-characterized BY and RM strains and generating a pool of BY/RM recombinant haploids (see Methods for details). Flow cytometry revealed that this pool encompassed three cell populations: non-fluorescent cells and, in roughly equal proportions, green and red cells (Figure 1D). We used fluorescence-activated cell sorting to isolate populations of green and red cells and then sequenced them in bulk to determine the BY and RM allele frequencies in each population. The BY parent strain contributes the MATa allele of the MAT locus, located on chromosome 3, while the MATα allele originates from the RM parent. As expected, the MATa allele was highly enriched in the green fluorescent recombinant haploids, while the MATα allele was enriched in the red population (Figure 1E). This result demonstrates that the fluorescent markers can be used to isolate populations of MATa, as well as MATα, haploids.

In addition to the expected large allele frequency differences at the MAT locus, we identified several allele frequency skews that were highly concordant between the two populations (Figure 1E). These skews correspond to previously identified growth QTL [17,18]. For example, the skew on chromosome 12 is located at the gene *HAP1*; the BY allele of this gene results in partial loss of function and poorer growth compared to the RM allele, leading to the observed reduction in frequency of the BY allele. The agreement between the allele frequency distributions in the two mating type pools indicates that the isolated cell populations can be used to map QTL with excellent reproducibility in both mating types.

### Phenotyping of wild strains and design of the round-robin cross

We set out to use our approach to dissect phenotypes mediated by different MAPK pathways. We used two stress conditions to query two different MAPK-signaling networks: high salt concentrations and caffeine. High salt conditions lead to activation of the HOG signaling cascade, while caffeine triggers the cell-wall integrity pathway.

We determined sodium chloride and caffeine tolerances for a collection of 65 *S. cerevisiae* isolates [24] and selected twelve diverse strains that spanned a range of resistances for our mapping studies (Figure 2A, Supplemental Table S1). These twelve strains vary at approximately 80% of the common SNPs (allele frequency ≥ 0.05) segregating within the larger strain collection. We designed a round-robin crossing scheme in which each of the twelve strains was crossed to two other strains, for a total of twelve crosses (Figure 2B). The round-robin crossing scheme is the most efficient way to intercross a set of strains while maintaining equal contributions of each strain [25]. Most mapping studies are based on crosses between resistant and sensitive strains, which implicitly assumes that resistance alleles are shared between resistant strains. By contrast, we randomly assigned the order of strains within the round-robin design without regard to strain phenotypes. As such, the crossing scheme included crosses between parent strains with different, as well as with similar, resistance phenotypes.

**Figure 2:**
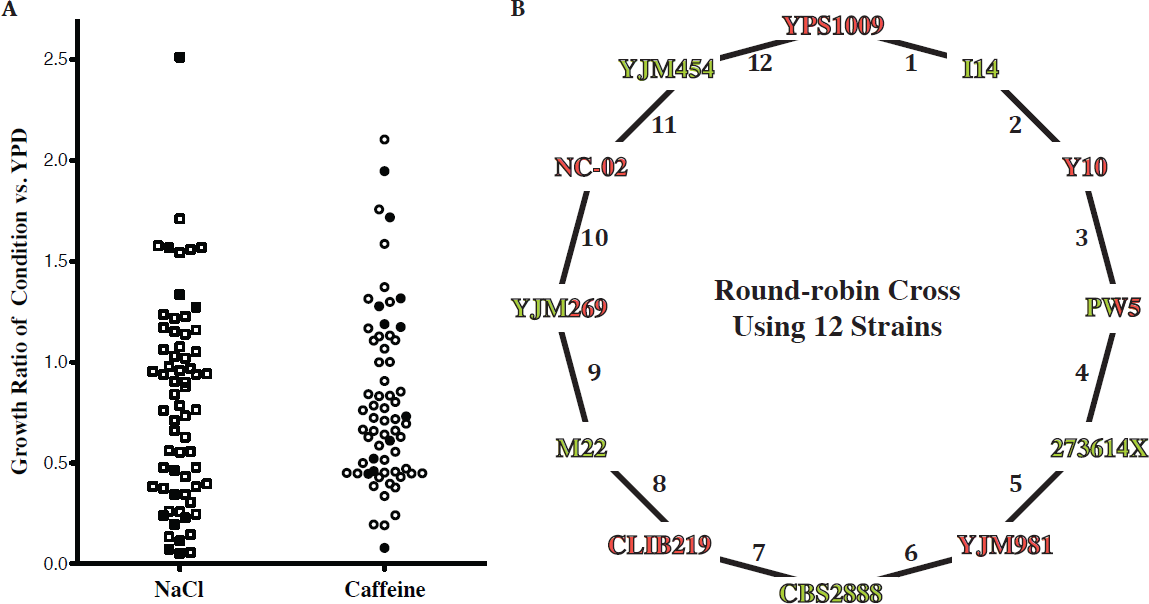
Design of round-robin cross to map salt and caffeine resistance. **(A)** Phenotyping of 65 diverse strains. Growth under the indicated condition (YPD plus 1M sodium chloride (NaCl) or 15 mM caffeine) was adjusted for growth under the permissive YPD condition (also see Supplemental Table S1). We picked strains (filled symbols) that are representative of the genetic and phenotypic diversity and crossed them according to a round-robin design **(B)**. Strains were crossed independent of phenotype and genetic relationship. Strain name colors indicate mating type (Green = MATa, Red = MATα)(also see Supplemental Figure S1).

To assess the genetic complexity of salt and caffeine resistance, we isolated up to twenty individual segregants from each cross. We measured growth of these segregants and the parent strains under high salt and caffeine stress conditions (Supplemental Figure S1). Three of the 24 phenotype distributions of the cross progeny were approximately bimodal, suggesting one large-effect determinant of resistance in these crosses. Other crosses exhibited phenotype distributions consistent with directional genetics and transgressive segregation [26]. In two crosses, the mean progeny phenotype was significantly lower than the midparent phenotype (Supplemental Table S2), indicative of interactions between the underlying causal loci [26,27]. These findings illustrate the genetic complexity of the resistance phenotypes among wild isolates.

### Mapping MAPK-dependent traits within the round-robin crosses

Next, we established a pipeline to map QTL between any pair of strains. Sequences of parent strains were used to generate lists of SNPs that segregated within a cross and could serve as genetic markers for QTL mapping. Mapping populations of both mating types were isolated and grown under permissive as well as selective conditions (YPD, YPD containing 0.5M sodium chloride, 1M sodium chloride, 15 mM caffeine or 20 mM caffeine). The resulting populations were bulk-sequenced to assess allele frequencies at SNP sites segregating in a particular cross. We used the MULTIPOOL software to calculate LOD scores testing whether allele frequencies at a given position in the genome in each selected population differed from those in the corresponding control population (Supplemental Figure S2)[19,28].

We used replicate BY/RM mapping experiments to determine the robustness and reproducibility of our mapping approach (see Methods for details). Allele frequency distributions of replicate experiments were extremely similar: comparisons between replicates did not exceed a LOD of 2.62 (Supplemental Figure S3 A & B). At a LOD score threshold of five, QTL detection was highly reproducible, with 90.1% of QTL reproducibly detected in replicate experiments (Supplemental Figure S3 C & D, Supplemental Table S3). Reproducibility was also high between selections based on mapping populations of opposite mating types (88.3%).

We applied our mapping strategy to identify QTL that influence MAPK-dependent resistance traits in the twelve Round-robin crosses. We used the MATa and MATα selections for each cross as replicate mapping experiments (see Supplemental Figure S4 for combined LOD plots for each cross). We used a stringent criterion that a QTL must exceed a LOD threshold of five in both selections, and identified 155 QTL across the twelve crosses and the different conditions. Of QTL identified in only one of the two matched experiments, the majority had LOD scores close to the LOD threshold (median LOD 6.7 for unreplicated QTL vs. 12.3 for replicated QTL, Supplemental Figure S5A). Indeed, for 93.5% of unreplicated QTL the direction of the underlying allele frequency skews was the same in the matched MATa and MATα experiments (permutation based *p* <0.001). This suggests that many of these QTL stem from small effect variants that narrowly escaped detection in one of the two experiments.

Next, we asked how QTL are distributed and shared among the Round-robin crosses (Figure 4 A & B, Supplemental Table S4). The salt QTL fell into 37 unique genomic regions, while the caffeine QTL comprised 23. There were 13 QTL regions that recurred between the conditions. We found an average of 6.2 salt resistance QTL (range 1 to 13) and 3.7 caffeine resistance QTL (range 0 to 7) per cross. Interestingly, the number of QTL detected per cross was independent of the genetic distance of the parent strains involved (divergence & salt QTL: Pearson correlation 0.154, divergence & caffeine QTL: 0.087). However, when the QTL of the two conditions were considered together per cross, this resulted in a weak correlation with the extent of parental divergence (0.257). We tested if the number of QTL per cross was a function of the phenotypic difference of the parent strains, since a previous study found that a higher number of QTL could be identified between yeast strains of similar phenotypes [29]. We found that this was the case for the salt resistance QTL (phenotypic difference & salt QTL: Pearson correlation −0.517), but not for caffeine resistance QTL (phenotypic difference & caffeine QTL: 0.395). While increasing the phenotypic similarity of yeast parent strains through the experimental fixation of large effect QTL can permit the identification of additional small effect QTL [30,31], this trend is condition-dependent for crosses between different wild isolates.

Because each genetic variant is introduced into the Round-robin scheme at least twice, each QTL is expected to be identified at least twice assuming 100% power and no genetic interactions. Contrary to this expectation, previous studies involving sets of interconnected crosses have found that the majority of QTL are ‘context-dependent’, that is their identification was dependent on a specific cross and not just on the parent strains involved [21,29,32]. Indeed, 33 of the 60 QTL regions we identified across the two conditions were found in only one cross (Supplemental Figure S5B). The overall lower LOD scores of these ‘context dependent’ QTL as a class only partially explained their lack of detection in additional crosses (Supplemental Figure S5C; Pearson correlation 0.417, *p* = 2.6 × 10^−6^). As such, the context dependency of these QTL is likely to be in part a result of interactions between the QTL and the genetic background [32].

### Functional connections of QTL to signaling

We asked if the QTL mapping had resulted in any functional enrichment in the underlying genes. While genes within caffeine resistance QTL were only marginally enriched for the GO-term ‘Response to Stimulus’ (p = 0.077), genes found under salt resistance QTL were significantly enriched for ‘Cellular Response to Stimulus’ (p = 0.006) and ‘Kinase Activity’ (p = 0.002) (Supplemental Table S5). This suggests that our mapping experiments point towards genetic variants that impact cellular signaling pathways.

Several of the identified QTL tower among the genetic architectures of the two resistance traits (Figure 3 A & B, Supplemental Table S4). Tolerance of high salt concentrations is dominated by variation at the *ENA* locus on chromosome 4, which we detected in eleven out of twelve crosses. The *ENA* genes encode sodium efflux pumps that, in the presence of high salt, are transcriptionally activated by the HOG pathway [33]. European *S. cerevisiae* isolates carry an *ENA* variant that resulted from an introgression from the yeast species *S. paradoxus* and exists in varying copy numbers, referred to as *ENA1* through *ENA5*. In contrast, non-European strains carry a single copy of the ancestral *ENA6* gene [34]. Both the introgressed variant and its copy number are associated with high salt tolerance [10]. We determined the copy numbers of the two *ENA* variants within our parent strains and found that strains with high salt resistance indeed carried multiple copies of the introgressed *ENA* gene (Supplemental Table S6). Interestingly, the identified *ENA* QTL reflected differences beyond the two divergent *ENA* variants and copy number. We identified *ENA* QTL in crosses between parents with equal copy numbers of the introgressed variant (Cross 9), as well as between non-European isolates carrying alleles of the *ENA6* gene that differed from each other by three missense mutations (Cross 12). Our findings corroborate the importance of the *ENA* locus in shaping the genetic architecture of salt resistance [10] and further illustrate the multi-allelic nature of this locus.

**Figure 3:**
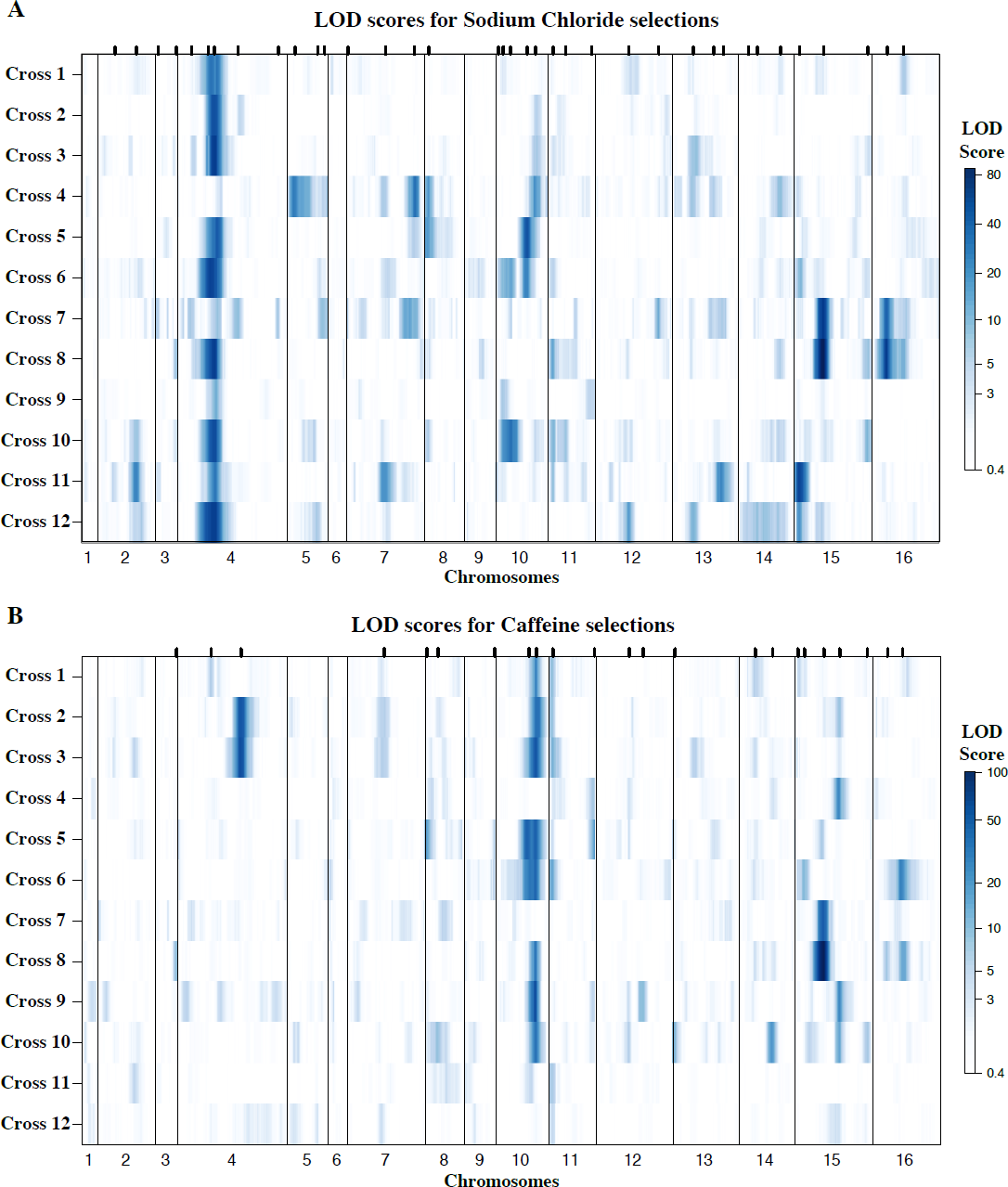
Genetic architectures of resistance identified in the round-robin crosses. Genome-wide LOD scores for **(A)** sodium chloride and **(B)** caffeine experiments are plotted jointly for the 12 Round Robin crosses to illustrate the how QTL are shared among the different crosses (see Supplemental Figure S4 for individual plots). The large effects of the multi allelic *ENA* locus on salt and *TOR1* on caffeine resistance are readily apparent. At each position the minimum of replicate selections is plotted for each cross. Tick marks on the upper axis indicate peak positions of QTL identified (QTL are listed in Supplemental Table S4).

The hallmarks of *S. cerevisiae* caffeine resistance architecture are the QTL containing *TOR1* and *TOR2* on chromosomes 10 and 11, respectively (Figure 3B). In contrast to other eukaryotes, yeast carries two copies of *TOR.* As in higher organisms, TORC complexes respond to nutrient signals to regulate a plethora of cellular processes through two functionally distinct complexes. In yeast, Tor1p is specific to TORC1 complexes, while Tor2p is able to participate in both TORC1 and TORC2 complexes [35]. Caffeine inhibits Tor1p function [36], leading to the activation of the CWI pathway [37]. We identified a *TOR1* QTL in eight of the twelve crosses, illustrating the large impact of *TOR1* on the genetic architecture of caffeine resistance. *TOR2* QTL were found in five crosses, and their identification likely speaks to Tor2p’s ability to functionally compensate for Tor1p in TORC1 complexes.

However, the QTL we detected as encompassing *TOR* genes were not mutually exclusive. In the four cases where we found both *TOR1* and *TOR2* QTL within the same cross, resistance alleles originated from the same or different parent strains an equal number of times. Furthermore, TOR negatively regulates transcription factors that are activated by the HOG pathway in response to stress (reviewed in [11]). Indeed, we uncovered salt resistance QTL containing *TOR2* in three crosses (Supplemental Table S4).

In addition to these QTL shared among many of the Round-robin crosses, several other QTL had large effects in a smaller number of crosses and contain genes with well-established roles in cellular signaling. A caffeine resistance QTL shared between crosses 2 and 3 encompasses *MSS4*, which encodes the essential Phosphatidylinositol-4-phosphate 5-kinase required for activation of the CWI MAPK pathway [38]. Two additional large effect QTL highlight the interconnectivity of the cellular signaling network. These QTL on chromosomes 10 and 15 strongly influenced growth in the presence of both high salt and caffeine. The QTL on chromosome 10 contains *CYR1*, the adenylate cyclase essential for cAMP production and as such cAMP-PKA signaling [39]. Similar to TOR, the cAMP-PKA pathway responds to nutrient signals to modulate a myriad of cellular processes, including the negative regulation of stress response transcription factors (reviewed in [11]). The QTL on chromosome 15, on the other hand, includes a gene that represents a functional counterpart to this aspect of cAMP-PKA signaling: *WHI2* encodes a phosphatase required for activation of the general stress response [40].

### Impact of strain history and source on QTL identified

While trait variation among *S. cerevisiae* isolates is great in comparison to related fungal species [10], the evidence for adaptive alleles is scant [9,41]. Rather than being strongly shaped by their source environment, yeast phenotypes have been proposed to be a result of population history [10]. The *ENA* locus illustrates this pattern. The introgression of the *S. paradoxus ENA* variant is widespread in European isolates regardless of their source environment. Our parent strains included European clinical and vineyard isolates, yet these ecological niches had no bearing on the *ENA* QTL we identified in crosses involving these strains. We wondered if alleles, denoted by the strain they derived from, tracked with source environment for any of the other QTL. We examined QTL detected in non-overlapping crosses (Supplemental Table S7), which excludes QTL likely due to single instances of loss- or gain-of-function mutations, and found two QTL where the grouping of alleles was unlikely to be a result of shared population history. Caffeine resistance alleles that gave rise to *TOR2* QTL belonged to wine and clinical strains from a diverse range of geographic sources, while the corresponding sensitive alleles stemmed from natural isolates, such as oak strains. A salt resistance QTL on chromosome 15, containing *YGK3*, which encodes a stress response controlling kinase, exhibited a similar pattern: a European and an American clinical strain contributed resistance alleles, while diverse natural isolates supplied sensitivity alleles. Further studies are needed to clarify whether these two cases are coincidental or whether they represent cases of environmental adaptation.

### Identification of MAT-dependent growth QTL

Yeast mating, akin to the stress responses examined above, is regulated by a MAPK signaling pathway. In addition to providing us with independent selection experiments, our ability to isolate mapping populations of either mating type allowed us to search for variants that influence growth through a genetic interaction with the mating type locus. Fitness differences between the two mating types have been reported [42,43], and we have shown that the MAT locus influences the differential expression of numerous genes [13]. In order to detect MAT-dependent growth QTL, we directly compared the allele frequencies resulting from corresponding MATa and MATα selection experiments.

For the round-robin crosses, we first compared the permissive YPD selections for each mating type. Among the twelve crosses, we identified six significant QTL (Supplemental Table S8). One of these QTL appeared in two non-neighboring crosses and encompassed the mating pathway regulator *GPA1* located on chromosome 8. Polymorphisms within Gpa1, which negatively regulates the mating pathway, can impair its function and hence result in costly activation of the pathway in the absence of mating pheromone [44]. Importantly, the *GPA1* allele selected for in one of our QTL experiments (Figure 4) contains a known fitness-associated variant [44,45]. We previously uncovered a genetic interaction between the MAT locus and *GPA1* that affects the expression of other mating pathway genes [13]. While this interaction did not result in a growth QTL in the BY/RM cross, our identification of MAT-dependent *GPA1* QTL suggests that this interaction can have an effect on growth in other genetic backgrounds. In addition to the *GPA1* QTL, we identified a QTL containing *SST2*, which encodes the GTPase-activating protein for Gpa1 [46].

**Figure 4:**
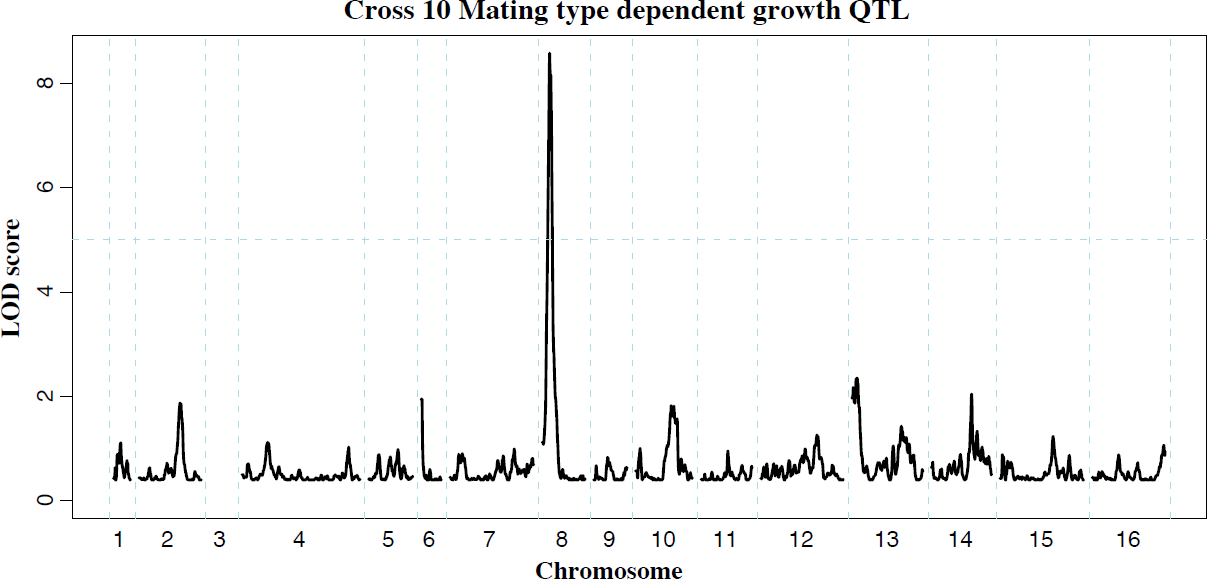
Mating type dependent QTL. Comparison of allele frequencies within MATa and MATα populations revealed loci that affect growth in a mating type dependent manner. The QTL on chromosome 8 encompasses *GPA1*, a negative regulator of the mating pathway that affects fitness in different strain backgrounds [1]. Chromosome 3 was omitted as it contains the mating type locus itself (see Supplemental Table S8 for all mating type dependent QTL identified).

Next, we asked if mating type had any influence on the allele frequency distributions arising under the two MAPK-inducing stress conditions. We found 19 instances of MAT-dependent stress resistance QTL (Supplemental Table S8). While it is difficult to determine whether these QTL represent an instance of MAT-dependence or failure to replicate a weak QTL, 14 of these QTL were identified in independent experiments using different dosages of the same stress condition or in both stress conditions. Among the latter group, we found that the *GPA1* and *SST2* mating type dependent QTL persisted in the presence of either stress condition. The mating and HOG MAPK pathways share the MAPKKK Ste11, and several studies have investigated how signaling specificity is achieved despite this shared signaling component [47,48]. In particular, faithful signaling along either pathway could be maintained by either mutual inhibition [47] or pathway insulation [48]. Our results suggest that the baseline fitness cost of the mating pathway [44] remains unaltered in the face of stressors that activate other MAPK cascades, which is consistent with the insulation of the mating MAPK from other MAPK signaling cascades [48].

### Utilizing the Round-robin design to identify causative variants

The Round-robin design enables a strategy to identify candidate causative variants that is not available using pairwise crosses. Specifically, the segregation of individual variants in relation to the presence or absence of QTL in multiparental crosses can be leveraged to reduce the search space of candidate causative variants [49,50].

Many sequence variants occur in several of the Round-robin crosses and, assuming additive effects independent of genetic background, should have similar consequences in these different crosses. We generated de novo assemblies of the parent strain genomes [51] and cataloged the non-synonymous coding variants in each QTL. We determined whether a given variant segregated within each cross by examining if the respective parent strains carried different alleles of the variant. For each QTL, we then assessed to what extent the segregation of a particular variant in each cross was associated with the detection of the QTL in the crosses (Supplemental Table S4). We used the maximum LOD scores within a particular QTL interval for each cross as a quantitative measure of QTL detection. This association analysis resulted in a 17-fold reduction in the number of variants to be considered as potentially causative for each QTL. We confirmed causality of individual sequence variants in two QTL to illustrate how this approach can aid in the identification of causal variants.

We first focused on the QTL on chromosome 15 that contained the stress response regulator *WHI2* [40] and strongly influenced the resistance to both salt and caffeine in crosses 7 and 8 (Figure 5A). Both crosses shared CLIB219 as a parent strain, and in both sets of experiments, the CLIB219 allele was strongly selected against, indicating that it leads to stress sensitivity. As this QTL did not appear in this form in other crosses besides the two involving CLIB219, the causative variant likely was private to CLIB219. Four variants out of 64 variants in the region were private to CLIB219 (Figure 5B). One of the variants was a frame-shift mutation in *WHI2*. Allele replacements of the CLIB219 allele of *WHI2* resulted in reduced growth in the presence of high salt and caffeine and thus confirmed that it was indeed the causal variant (Figure 5C).

**Figure 5:**
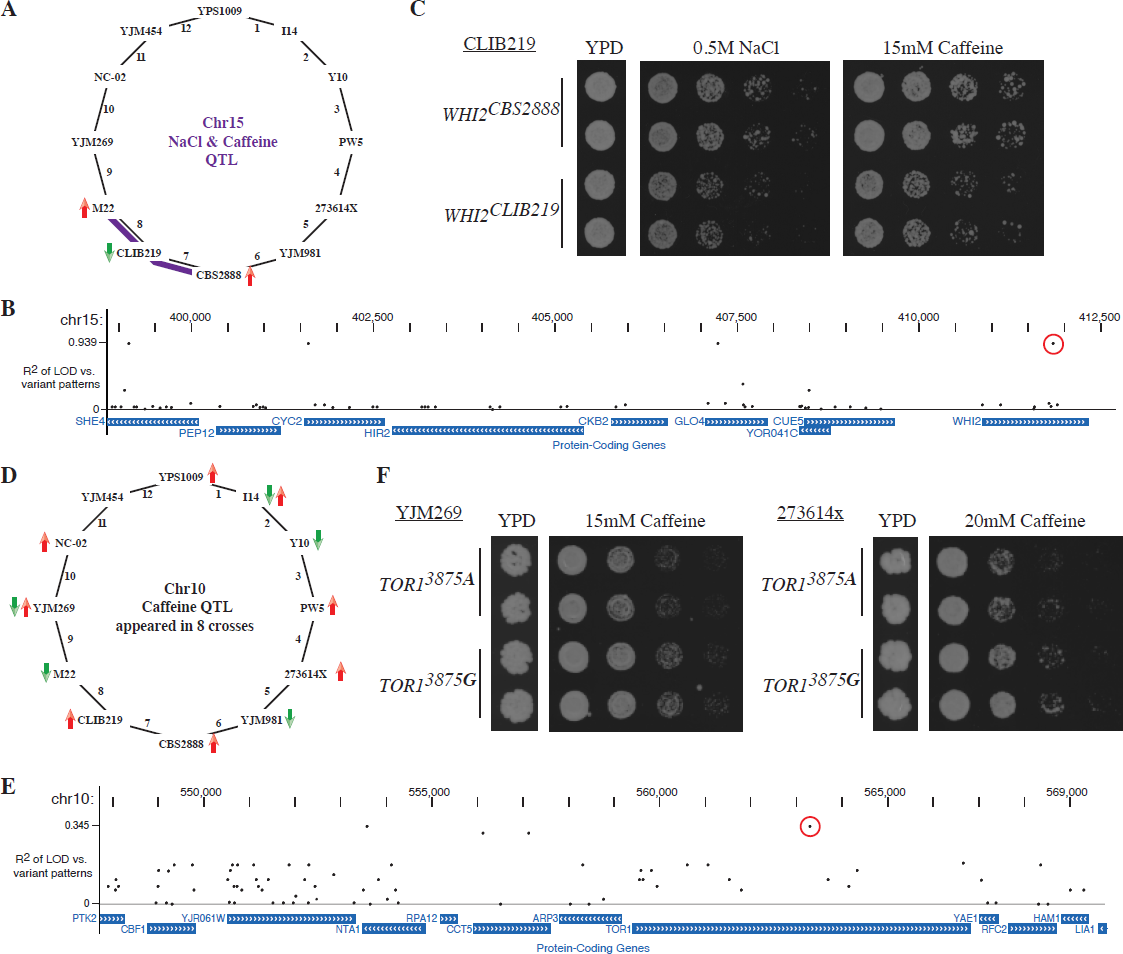
Leveraging the Round Robin design to identify causative variants. **(A)** A QTL on chromsome 15 affects salt and caffeine resistance in crosses 7 and 8 (with smaller effects in other crosses). The CLIB219 is shared between the two crosses and in both cases it contributes the allele is selected against, suggesting that the causative variant is private to CLIB219. **(B)** We determined the association of coding variants within this QTL interval with the observed pattern of LOD scores. The highest association scores were exhibited by four variants private to CLIB219; one a frame-shift mutation in *WHI2.* **(C)** Allele replacements of *WHI2* in the CLIB219 strain background. The CLIB219-specific frame shift mutation resulted in decreased growth in the presence of both high salt and caffeine. **(D)** Identification of a quantitative trait nucleotide (QTN) within the caffeine resistance QTL on chromosome 10. The QTL was identified in eight crosses and is multi-allelic as specific alleles were beneficial or deleterious depending on a particular cross. **(E)** Measuring the association of variants and LOD scores confirmed that no single variant could explain the observed pattern of QTL and indicated a *TOR1* SNP (3875 G −> A) as a QTN (red circle). **(F)** Introduction of the *TOR1^3875**A**^* variant into two strain backgrounds, YJM269 and 273614x, reduced resistance to caffeine. For each allele replacements two independent transformants are shown.

Next, we turned to a QTL with a substantially more complex allelic pattern. The *TOR1*-containing caffeine resistance QTL on chromosome 8 was identified in eight crosses. Specific alleles were selected for or against depending on the particular cross, which was a clear indication that multiple alleles were present at this QTL (Figure 5D). While no single variant perfectly recapitulated the pattern of LOD scores (Figure 5E), two SNPs in *TOR1* and *NTA1* scored the highest. Closer examination of the segregation pattern of these SNPs revealed that they were common to two caffeine sensitive strains, M22 and YJM981. Allele replacements of the *TOR1^3875**A**^* variant in two strain backgrounds confirmed that it contributes to caffeine sensitivity (Figure 5F). These results demonstrate how the Round-robin crossing scheme can be leveraged to focus the list of candidate causative variants.

## Discussion

Here, we have demonstrated the ability of a novel QTL mapping approach to query a large proportion of *S. cerevisiae* genetic diversity. We employed this approach to characterize the genetic architectures of three MAPK-dependent traits: resistance to high salt or caffeine and growth differences due to mating type. The relationship of the identified genetic variants to MAPK signaling pathways ranged from upstream modulators to core regulators and downstream targets.

Our modified X-QTL approach makes it possible to rapidly generate large mapping populations without extensive strain engineering. Quantitative trait mapping studies in yeast have uncovered the genetic architectures of traits central to biology, such as gene expression and protein-level variation [14,19,52], as well as traits of medical and industrial importance [15,53]. Yet, these studies have only scratched the surface of the phenotypic variation of *S. cerevisiae* strains [10,20], a space that is sure to expand with the continued discovery of additional yeast isolates [54,55]. Our method promises to be of great utility in future yeast QTL mapping studies. Here, we have chiefly used our approach to isolate large pools of segregants for BSA-based QTL mapping, but we also illustrate its utility in isolating individual segregants that can be used in traditional linkage-based QTL mapping studies. In contrast to previous methods [18,56], our method can be employed in nearly any strain background with minimal strain engineering and enables positive selection for haploids of either mating type. Furthermore, the fluorescent mating type markers are amenable to combination with fluorescent readouts of cell physiology or gene expression [18,19]. Such reporter combinations will make it feasible to isolate mapping populations and select for phenotypes of interest simultaneously.

We have genetically dissected stress resistances mediated by two MAPK pathways, as well as interrogated the genetic contribution to the baseline cost of the mating pathway. We studied twelve diverse yeast strains that represent a large proportion of the genetic and phenotypic diversity of *S. cerevisiae*. Thus the QTL identified capture much of the species-wide genetic variation that influences the phenotypic consequences of these MAPK pathways. We identified QTL that encompass genes connected to MAPK signaling at multiple levels (Supplemental Table S4). Several QTL contained genes whose protein products are responsible for the first step of the MAPK signaling cascades: osmosensors (*MSB2* & *SHO1*) and CWI sensors (*WSC3* & *MTL1*) responsible for detecting the respective perturbations. Moving along the signaling cascades, we found several key regulators such as *MSS4* and *GPA1*. Furthermore, one of the caffeine tolerance QTL contained *SLT2*, which encodes the MAPK at the center of the CWI response. Arriving at the output of the signaling pathways, we identified several genes transcriptionally activated in response to stress. In addition to the *ENA* genes, we found *GPG1*, which is activated by the HOG pathway to increase glycerol synthesis to counteract changes in osmolarity [57]. Finally, although the resistance phenotypes of the parent strains were not correlated, thirteen QTL occurred under both conditions. The overlap between QTL is consistent with crosstalk between the two stress responsive pathways [58,59].

In addition to genes directly associated with MAPK pathways, we found genes that highlight the interconnectivity of the cellular signaling network. Variants in *CYR1* and *TOR2*, core members of the cAMP-PKA and TOR pathways respectively, were tied to both stress responses. These pathways sense the internal availability of nutrients and repress the general stress response [11]. The TOR pathway in particular regulates a myriad of functions that mediate cell growth [60]. In contrast to higher eukaryotes, yeast carries two copies of the *TOR* gene. Our study links numerous alleles of *TOR1* and *TOR2* to differential growth under stress conditions.

The two Tor proteins are redundant in regards to their participation in TORC1 complexes, but only Tor2p can partake in TORC2 complexes [35]. Despite this difference, the two *TOR* genes are diverging at the same rate as the majority of duplicated gene pairs [61]. However, these studies were based on individual strains. Our QTL results point towards functional variation in both Tor isoforms and raise questions about how they jointly regulate cell growth in different yeast isolates. Are the roles of the two proteins in TORC1 complexes truly equivalent, or are the *TOR* alleles indicative of functional tradeoffs? Similarly, it bears determination whether stress sensitive alleles of the *TOR2* QTL, which were found in natural rather than ‘domesticated’ isolates, are beneficial under conditions approximating their source environment.

The identification of *WHI2* further echoes the interplay between internal nutrient and external stress sensing pathways. While *WHI2* is required for the activation of stress response pathways [40], its loss leads to prolonged growth under nutrient limited conditions [62]. Interestingly, *WHI2* loss-of-function alleles were found to be driver mutations in experimental evolution experiments [63,64]. Variants that abrogate Whi2 function were also discovered as recurrent secondary site mutations in yeast strains with specific gene deletions [65]. As such, *whi2* mutants not only illustrate how laboratory evolution experiments can recapitulate the biology of wild isolates, but also suggest that the concomitant loss of stress responsiveness is adaptive under permissive growth conditions [63,64]. Indeed, the *WHI2* allele responsible for stress sensitivity was advantageous for growth under the permissive YPD condition.

The QTL results provide insights into the types of genes that carry variants shaping MAPK-associated phenotypes of different yeast isolates. The core elements of the HOG and CWI signaling cascades (HOG: *SSK1*, *SSK2*, *STE11*, *PBS2*, and *HOG1*; CWI: *RHO1*, *PKC1*, *BCK1*, *MKK1*, *MKK2,* and *SLT2*) are highly conserved across species. Analysis of the strain sequences revealed that despite their species-level conservation, all of these core proteins, with the exception of Rho1p, contain instances of non-synonymous coding variants. Yet, we only identified QTL at *SLT2*, indicating that the coding variants in the other core genes may be of little functional consequence.

In contrast to the central cascade, proteins responsible for sensing perturbations in osmolarity or cell wall integrity diverge rapidly between yeast and related fungal species [12,66]. Divergent sensor proteins in other species are thought to enable the response to different habitats. For instance, other fungi use the HOG pathway to respond to a broader range of stresses than *S. cerevisiae* (reviewed in [11]). Consequently, we wondered if variation in sensor genes underlies the observed stress resistance variation among different *S. cerevisiae* isolates. We found stress sensors at two QTL for both salt and caffeine resistance, but these QTL had low LOD scores indicative of small effect variants. This is likely explained by the large degree of redundancy among the sensor proteins [67,68]. Osmosensors are encoded by a set of five genes, and CWI sensors are encoded by six genes. While sensor redundancy allows for species-level fine-tuning of the stress responses, any single change is unlikely to have large phenotypic consequences. The same holds true for the large number of transcriptional targets of the MAPK pathways, with the exception of special effectors such as the introgressed *ENA* variants and *GPG1* [69]. In contrast, many genes associated with large effects, such as *CYR1, GPA1, MSS4* and *TOR1/2*, have functions essential to the MAPK and other signaling pathways. Together, these findings illustrate how genetic variants shape the phenotypic space of MAPK stress resistance traits within *S. cerevisiae*. The global picture that emerges from our results is that genes thought to underlie phenotypic differences between fungal species due to their high levels of divergence contribute little to trait variation among *S. cerevisiae* isolates.

The QTL we identified fall into several different classes in regards to their appearance among the twelve round-robin crosses. 33 of the 60 grouped QTL were ‘context dependent’, meaning they were detected in a single cross. The preponderance of ‘context dependent’ QTL was also notable in studies involving smaller panels of pairwise crosses [21,29]. Context-dependency has been attributed to interactions between QTL and the genetic background. In particular, large effect variants can mask the presence of small effect variants [30,31]. For salt resistance, the negative correlation between parent strain phenotype differences and the number of QTL identified was largely driven by the large effect of the *ENA* locus. For example, Cross 7 between two sensitive parent strains lacking the *ENA* introgression permitted the detection of a large number of small effect QTL. Of the twelve QTL detected in exactly two crosses, seven appeared in crosses involving the same parent strain, and as such can be classified as ‘strain dependent’ [29]. This set of QTL likely results from single alleles private to the shared parent strain [21]. Beyond this simple case, deducing the number of underlying alleles and their frequency becomes difficult. For instance, of the eleven QTL appearing in three crosses, nine were found in a pair of neighboring crosses plus one additional cross. Several models could explain this pattern, and we attempted to use the segregation pattern of the sequence variants within each QTL to determine which alleles best explained the observed QTL pattern. While this approach resulted in a significant reduction in the number of variants to consider, more than twelve crosses will be necessary to distinguish between the possible allele patterns with certainty. Going forward, our fluorescent markers allow facile creation of segregant pools from many additional crosses, as well as large mapping panels of individual segregants [17]. In conjunction with new genome engineering technologies, these approaches will enable even more complete genetic dissection of MAPK signaling and other complex traits.

## Acknowledgements

We thank members of the Kruglyak lab and Jennifer Leslie for helpful discussions and comments on the manuscript. We thank Arthur Shockley for technical help. We are grateful to members of the Botstein lab for various strains and reagents. We are especially thankful to Christina DeCoste in the Princeton Flow Cytometry Resource Facility for outstanding technical assistance and experimental advice. This research was supported by National Institutes of Health (NIH) grant R01 GM102308, a James S. McDonnell Centennial Fellowship, and the Howard Hughes Medical Institute (LK), a NIH postdoctoral fellowship F32 GM101857-02 (ST), a research fellowship from the German Science Foundation AL 1525/1-1 (FWA), and a National Science Foundation (NSF) fellowship (JSB).

## Supplemental Figure Legends

<u>Supplemental Figure 1: Phenotyping of individual Round Robin segregants</u>

Between 17 and 20 recombinant haploids were isolated for each of the twelve round robin crosses and, in conjunction with the haploid parent strains, phenotyped as to their growth in the presence of **(A)** 1M NaCl and **(B)** 20 mM Caffeine. Parent strains were phenotyped using five biological replicates. Segregants and parents of crosses 4, 7 and 9 were tested separately from those of the other crosses; hence the phenotypes of shared parent strains are not perfectly matched. Growth on the restrictive conditions was normalized using growth under the permissive YPD condition.

<u>Supplemental Figure 2: Conversion of allele frequencies to LOD scores</u>

**(A)** Allele frequency plots for a BY/RM MATa caffeine selection and its corresponding YPD control experiment. Loess-smoothed BY allele frequencies across the genome are plotted. **(B)** To account for growth effects under the permissive YPD growth condition, allele frequencies distributions resulting from growth on YPD are subtracted from those produced by the selective condition. Grey points represent individual alleles at which allele frequencies were measured and illustrate the dense coverage of these genetic markers. **(C)** Allele frequencies are converted into LOD scores using the MULTIPOOL software [2]. Following the same principle as subtraction of control from selection allele frequencies, the MULTIPOOL software calculates LOD scores based on the differences between the two allele frequency distributions. QTL passing the LOD threshold of 5 (dashed horizontal blue line) are demarked on the upper axis and their 2-LOD mapping intervals indicated by orange vertical bars.

<u>Supplemental Figure 3: Allele frequency and QTL plots of replicate BY/RM selection experiments</u>

Genome-wide allele frequency distributions were determined in eight replicate selection experiments each for **(A)** salt and **(B)** caffeine. The experiments were carried out in duplicate for each of two independent transformants of the BYxRM diploid with the mating type marker construct. MATa and MATα selections experiments for each provide for further replication, resulting in a set of eight replicates for each condition. Lines represent moving averages of the allele frequency spectra and are overlaid to illustrate their reproducibility. LOD score plots of replicate **(C)** sodium chloride and **(D)** caffeine selection experiments illustrate the high reproducibility of our QTL mapping approach. Each selection was carried out in quadruplicate for each mating type. The dashed horizontal line indicates the LOD threshold of 5. Tick marks on the upper axis indicate peak positions of QTL identified.

<u>Supplemental Figure 4: Individual LOD plots for each of the round-robin selection experiments</u>

LOD plots of the MATa (Green) and MATα (Red) selection experiments for each cross and condition are jointly plotted. The dashed horizontal line indicates the LOD threshold of 5. The 2-LOD confidence intervals of the QTL identified are indicated using vertical orange bars. For Cross 4 under the 1M sodium chloride selection was very restrictive resulting in a low number of individual segregants in the genotyping pool and hence less reproducible LOD scores. Yet, the QTL confirmed by the replicate selection experiments were also detected using a less restrictive selection condition of 0.5M sodium chloride and largely (7 of 9 QTL) shared with other crosses.

<u>Supplemental Figure 5: Analyses of QTL identified in the round-robin cross</u>

**(A)** LOD score distribution of QTL detected in both of the MATa and MATα selections experiments compared to that of QTL only found in either of the two experiments. The majority of QTL that miss replication have LOD scores close to the threshold used to call QTL. **(B)** Histogram of the number of crosses per QTL group. Theoretically, assuming purely additive effects and perfect detection, each QTL should be observed in at least two crosses. **(C)** LOD scores of the QTL belonging to different classes of grouped QTL. Identification of QTL in only one cross is only partially explained by the lower LOD scores of this class of QTL (pearson correlation 0.417).

## Material and Methods

### Plasmid construction

Plasmids with drug markers were constructed in two steps. We first generated pRS416 and pRS426 Gateway expression vectors suitable for one and three fragment Gateway cloning. As described previously, the pRS vectors were digested with Sma1 and the Gateway cassettes introduced by ligation [1]. The drug markers were then introduced from a set of plasmids kindly provided by Mikko Taipale and Susan Lindquist (Whitehead Institute, Cambridge, MA). These Gateway drug markers plasmids were digested with BglII and XhoI in order to create fragments with the MX drug resistance markers alone. PCR was used to generate fragments of the pRS416 and pRS426 Gateway vector backbones with Xho1 and BglII cut sites and without the *URA3* marker. PCR products were gel extracted and digest with Xho1 and BglII. The vector backbone with the two different Gatewat cassettes and the three different drug resistance markers where then combined by ligation.

The *STE2* and *STE3* promoters were cloned upstream of the genes encoding the fluorophores mCitrine and mCherry using overlap extension PCR. Promoter fragments based on Tong et al. [2] were PCR amplified from genomic DNA, while fluorophores were amplified from plasmids kindly provided by Gregory Lang (Lewis-Sigler Institute, Princeton University, Princeton, NJ). The resulting fusions, as well as the *CYC1* terminator, were cloned into different Gateway entry vectors to facilitate their final combination in the three fragment Gateway plasmids.

Plasmids created for this study are listed in Supplemental Table S9, while primers used are listed in Supplemental Table S10.

### Strain construction

The fluorescent mating type markers were tested in BY strains kindly provided by the Botstein lab (BY 12045 MATa Δ*ura3*, BY 12046 MATα Δ*ura3*). The BY/RM diploid was published previously (yLK1993 [3]). Other strains were previously described in Schacherer *et al.* [4]. To facilitate the generation of diploids for the round-robin cross, the KanMX cassette used to delete *HO* was replaced by the HphMX cassette in MATx strains [5]. MATa and MATx parent strains were mated in YPD and subsequently streaked to YPD + G418 + Hygromycin to select for diploids. Putative diploids were tested for their ability to sporulate and to determine their ploidy [6]. The p41Nat plasmid carrying the fluorescent mating type markers was introduced by standard yeast transformation [7].

We used the NatMX cassette from pUG74 [8] to delete *URA3* gene in the haploid round-robin parent strains in order to make them amenable to allele replacement protocols. Attempts to make allele replacements according to Erdeniz *et al.* were unsuccessful [9]. We modified the approach described by Stuckey *et al.* to make allele replacements [10]. For the *WHI2* allele replacements, the entire *WHI2* ORF was first deleted using the pCORE construct [11] and then replaced by transformation with PCR-amplified *WHI2* ORFs plus approximately 250 bases up- and downstream from the different strains. For the *TOR1* allele replacements, we first integrated *K. lactis URA3* right at position 3875 of *TOR1.* We then removed the *K. lactis URA3* integration according to Storici *et al.* using complementary oligos spanning the integration site and carrying the two different *TOR1* alleles (*TOR1* 3875G and *TOR1* 3875A). Primers used in allele replacement experiments are listed in Supplemental Table S10.

### Strain phenotyping

We phenotyped a set of strains previously described by Schacherer *et al.* [4], as well as sets of random segregants stemming from the round-robin crosses. The strains were grown overnight at 30°C in 96-well plates containing YPD media according to standard yeast culture protocols [6]. The strains were then pinned in quadruplicate onto agar plates using a Singer RoToR HDA. Strains were phenotyped on YPD plates (the permissive control condition), as well as YPD plates containing 0.5M sodium chloride, 1M sodium chloride, 15 mM caffeine or 20 mM caffeine. Both sodium chloride and caffeine were autoclaved along with the other ingredients needed to make standard YPD plates. After strain pinning, plates were grown at 30°C in large plastic containers to avoid excessive evaporation. YPD plates were scanned after two days of growth, while stress condition plates were scanned once growth reached a level equivalent to that observed on the YPD plates [3]. Images of the plates were processed and analyzed to extract strain growth information as previously described [3]. Growth under the stress conditions was adjusted for growth under the permissive stress condition by calculating the ratio of the two growth measurements. We used a modified version of the epistasis test described by Lynch and Walsh [12] as detailed in Brem and Kruglyak [13].

### Spore isolation

We modified a published random spore prep protocol [6] in order to enrich for spores prior to FACS. Diploids were grown in 3 ml YPD overnight at 30°C. Cultures were spun down and washed with water. Cell pellets were resuspended in 1 ml sporulation media containing 25 mg/L ClonNAT or 200 mg/ml G418 to maintain the selection for fluorescent mating type marker plasmids. Half of the cell mixture was then added to 2.5 ml drug containing sporulation media. Sporulation cultures were incubated for four to six days at room temperature depending on the sporulation efficiency of a particular cross.

250 µl of the sporulation cultures was spun down for one minute in an eppendorf tube. After removal of supernatant the cell pellets were resuspended in 100 µl water. 17 µl of the cell suspensions was then added to new eppendorf tubes containing 3 µl of β-glucoronidase. After a brief vortex tubes were incubated in an Eppendorf Thermomixer at 30°C for two hours while shaking at 900 rpm. Digestion and opening of ascii after this two hour incubation was confirmed by light microscopy. Digestions were quenched by the addition of 100 µl water. In order to break apart digested ascii, tubes were vortexed for two minutes and then spun down for one minute. The supernatant was removed, another 100 µl was added and the tubes were vortexed for another two minutes. As hydrophobic spores stick to the sides of the eppendorf tubes all liquid is removed after the second vortex. Tubes were washed three times by addition of 1 ml water and inverting the tubes fully three times. After the washes, 1 ml 0.01% NP-40 (igepal) was added added to the tubes. Tubes were the sonicated using a tip sonicator for 1 min at power level 2.

In order to isolate individual spores or to assess the amount of spores isolated, dilutions of the spore prep (for most crosses 25 µl spore mix in 250 µl H2O was used) on the plates containing 25 mg/L ClonNAT or 200 mg/ml G418. Individual haploids were picked after two days of growth at 30°C using a fluorescence-equipped dissection microscope. For cell sorting 250 µl of the samples was plated on four plates containing the appropriate drug to select for the marker plasmid.

### Selection and FACS

Spore preps were harvested after 2 days of growth at 30°C. Cells were washed off plates using 5 ml water and collected in 50 ml conical tubes. Cells were sonicated for 1 min at power level 2 and then diluted to an OD of 0.4 for FACS in 1xPBS. We used a BD FACSVantage SE to sort 500,000 green and red fluorescent cells each into vials containing 2 ml YPD with 100 µg/ml ampicilin to prevent bacterial contamination during the cell sorting. The sorted cells were added to 10 ml YPD containing 100 µg/ml ampicilin in glass tubes and cultured for 6 hours at 30°C. Cultures were spun down for 5 minutes at 3,000 rpm and after removal of supernatant cells were resuspended in 950 µl water. 200 µl of cells suspension were then plated on 5 different conditions: YPD, YPD with 0.5M NaCl, YPD with 1M NaCl, YPD with 15 mM caffeine, and YPD with 20 mM caffeine. Plates were grown for 2 days at 30°C. Plates were made by adding the appropriate amount of NaCl or caffeine to the standard YPD recipe prior to autoclaving.

### DNA isolation and library prep

We assessed to what extent segregant populations grew under the selective conditions and proceeded with conditions that resulted in a reduced yet adequate amount of growth compared to the permissive YPD condition [14]. Cells were harvested by washing plates with 5 ml water and collected in 15 ml conical tubes. Cells were spun down and stored at −80°C after removal of supernatant. DNA was isolated from the harvested cells according to Lee *et al.* [15]. Cell pellets of approximately 50 µl were used for each DNA prep. DNA was isolated from parent strains in the same manner except that parent strains were grown overnight in 3 ml YPD at 30°C.

Sequencing libraries were generated using Epicentre Nextera DNA Sample Prep kits or Illumina Nextera Sample Prepartation kits as described previously [3]. Residual salts and other contaminants can cause Nextera library preps to fail. We cleaned the input DNA using the Zymo Research Genomic DNA Clean & Concentrator kits if the initial library prep failed. Sample libraries were indexed using primers designed according to Meyer and Kircher [16] and sequenced in groups of 25 per lane on an Illumina HiSeq sequencer.

### Allele frequency measurements and QTL mapping

We created a set of genetic markers for each cross by aligning parent strain sequences to the sacCer3 reference genome and we then used custom scripts written in the programming language R to identify SNPs that differed between any set of parent strains. We then determined allele frequencies at these variant sites. We subtracted the YPD allele frequencies at each variant position from those obtained under the different growth conditions to determine the allele frequency skews that are due to a specific condition rather than permissive growth. To determine allele frequency differences due to mating type, allele frequencies from MATα samples were subtracted from those of corresponding MATa samples.

We converted allele frequency skews into LOD scores using the MULTIPOOL method [17] with the following parameters: −t, −n 1000, −m contrast, −c 2200, −r 100. MULTIPOOL outputs were used to generate genome-wide LOD score plots [18]. LOD plots were generated in R. All the subsequent analyses were conducted using the statistical programming language R. We called QTL at positions that exceeded a LOD threshold of 5 and QTL regions using LOD drop intervals of 2, as described previously [18]. In order to combine QTL across replicate experiments we determined which QTL had overlapping mapping intervals. We then used the mean of replicated QTL intervals to identify QTL that overlapped between different round-robin crosses. We combined QTL that were detected at different concentrations of the same conditions (0.5M NaCl & 1M NaCl, 15 mM Caffeine & 20 mM Caffeine) to provide a conservative estimate of the QTL LOD scores and regions.

We used a set of replicate BY/RM mapping experiments to determine the false discovery rate and reproducibility of our mapping approach. We generated allele frequency measurements for four independent experiments using two independent transformants of the BY/RM diploid with the mating type marker plasmid. We isolated mapping populations of either mating type and used the same selection conditions as for the round-robin crosses (YPD, +0.5M NaCl, +1M NaCl, +15 mM Caffeine, +20 mM Caffeine). The YPD 20 mM Caffeine condition resulted in very low growth of BY/RM segregants and we did not proceed further with this condition. Given the two mating type mapping populations and four independent experiments, we generated eight selections for each condition. We compared the allele frequencies resulting from technical replicates (16 comparisons) stemming from one transformant and found no allele frequency fluctuations that resulted in LOD scores higher than 2.62. Analysis of biological replicates, i.e. replicates based on different transformants, resulted in six peaks with LOD scores higher than our threshold of LOD 5. Yet, five of these peaks were actually the same region identified multiple times in different replicate pairs suggesting that they were due to a genetic difference between the two BY/RM transformants and not due to random allele fluctuations. The remaining peak tagged the mating type locus on chromosome 3 pointing towards a difference in the mating type selection rather than a random allele fluctuation.

In order to asses the reproducibility of the mapping approach we determined condition-specific QTL for individual experiments and then asked to what extent QTL where found in pairs of replicates. To keep this assessment of reproducibility consistent with the round-robin mapping experiment, we did not ask whether QTL were identified multiple times across all replicates. Such a measure of reproducibility would be inflated in comparison to experiments with only two replicates, such as the experiments with the round-robin crosses. Rather we tested if QTL were reproducibly identified in designated pairs of experiments, either technical replicates performed with the same strain and the same condition or the experiments of opposite mating types of one replicate. At a LOD threshold of 5, 90.1% of QTL were reproducibly identified between technical replicates of the same mapping experiment. Reproducibility was 88.3% using mapping populations of opposite mating type. For the experiments with technical replicates, the directions of the allele effects of the QTL identified in only one experiment were consistent among replicates in 100% of the cases. This suggests that these QTL only narrowly missed the LOD threshold in replicate experiments. Indeed, their LOD scores were close to the threshold of LOD 5 (median LOD 5.9 vs. 9.84 for replicated QTL). For the round-robin experiments across all conditions, 74.4% of QTL (311 of 418 individually identified QTL, with one instance of one peak overlapping two in the second dataset, resulting in 155 jointly identified QTL) were identified in both of the mapping experiments. Since QTL that narrowly miss the LOD threshold are classified as not being reproducible, we asked to what extent allele frequency directions, in the control case BY versus RM, agreed over the identified region. We determined the mean allele frequencies for the region underlying a QTL and compared them across the pairs of replicate experiments. For the BY/RM experiments, the sign of the allele frequencies agreed for all eleven QTL that had missed replication. In the case of the round-robin crosses, allele frequency directions agreed for 93.5% of non-replicated QTL. This result was robust to permutation of the allele frequencies (p <0.001).

### Testing for association between variants and LOD scores

We used Cortex [19] to generate reference-assisted genome assemblies of the round-robin parent strains. We then used VCFtools [20] to subset the VCF file generated by Cortex to the individual regions of the QTL. We used VEP [21] to identify non-synonymous coding sequence variants within QTL intervals. We then generated a table for each QTL in the statistical programming language R that contained the replicate-averaged maximum LOD scores for each cross for the given QTL region and notations how each variant segregated among the crosses. For each variant, we then calculated the squared correlation between its segregation pattern and the pattern of LOD scores. We determined the distribution of for all the variants of a given region and used a p value of 0.01 to call significant variants. The coefficients of determination were visualized along QTL intervals using the UCSC Genome Browser [22].

